# MYO1C facilitates arrival at the Golgi apparatus through stabilization of branched actin

**DOI:** 10.1101/409110

**Authors:** Anahi Capmany, Azumi Yoshimura, Rachid Kerdous, Aurianne Lescure, Elaine Del Nery, Evelyne Coudrier, Bruno Goud, Kristine Schauer

## Abstract

We aim at the identification of myosin motor proteins that control trafficking at the Golgi apparatus. In addition to the known Golgi-associated myosins MYO6, MYO18A and MYH9 (myosin IIA), we identify MYO1C as a novel player at the Golgi. We demonstrate that depletion of MYO1C induces Golgi apparatus fragmentation and decompaction. MYO1C accumulates at dynamic structures around the Golgi apparatus that colocalize with Golgi-associated actin dots. Interestingly, MYO1C depletion leads to loss of cellular F-actin, and Golgi apparatus decompaction is also observed after the inhibition or loss of the Arp2/3 complex. We show that the functional consequences of MYO1C depletion is a delay in the arrival of incoming transport carriers, both from the anterograde and retrograde routes. We propose that MYO1C stabilizes branched actin at the Golgi apparatus that facilitates the arrival of incoming transport at the Golgi.

## Introduction

The Golgi apparatus is the main hub of intracellular trafficking, at the interface between anterograde and retrograde routes. Golgi membrane dynamics are mandatory for efficient sorting of cargos traversing the Golgi apparatus and require tensile forces from the cytoskeleton. Particularly, actin-dependent myosin motor proteins have been implicated in the formation of tubular membrane carriers and their fission from the Golgi: Historically, the non-muscle myosin IIA (MYH9) has first been proposed to function in the production of vesicles from the Golgi apparatus (Müsch et al., 1997). More recently, the depletion and inhibition of myosin II induced the appearance of long Golgi-derived, Rab6-positive, membrane tubules further supporting its role in fission events during *trans*-<u>G</u>olgi <u>n</u>etwork (TGN) exit (Miserey-Lenkei et al., 2010). Myosin VI family member MYO6, a myosin moving towards the minus ends of F-actin, has also been found on Golgi membranes and was shown to play a role in Golgi ribbon formation and exocytic events (Warner et al., 2003). Moreover, MYO18A, MYO1B and MYO5B were additionally proposed to control the formation of tubular and vesicular membrane carriers from Golgi and TGN membranes (Dippold et al., 2010; Almeida et al., 2011; Liu et al., 2013). Interestingly, although Golgi-associated myosins belong to different classes, they all have been proposed to facilitate the exit from the Golgi apparatus. Here we aimed at the systematic analysis of myosins that regulate trafficking at the Golgi level and the characterization of the underlying molecular mechanisms by which they regulate Golgi membrane dynamics.

## Results

### Identification of myosins that alter Golgi morphology

Defects in Golgi trafficking are characterized by changes in Golgi morphology (Goud et al., 2018). Thus, we systematically depleted 36 myosins present in human and sought for those which altered Golgi morphology. We employed our approach combining cell culture on adhesive micropatterns and quantification of organelle positioning by probabilistic density mapping (Schauer et al., 2010; Duong et al., 2012; Schauer et al., 2014). We have previously shown that when seeded on micropatterns, the Golgi apparatus has a characteristic, well-defined, stable and reproducible positioning allowing automated detection of subtle changes in its organization in a reduced number of cells (Schauer et al., 2010). In three independent experiments, immortalized human hTertRPE-1 cells were transfected with a siRNA library targeting each myosin with four independent siRNAs (Figure 1A). The Golgi apparatus was analyzed in fixed, crossbow-shaped micropatterned cells using our density-based statistical analysis as previously reported (Duong et al., 2012; Schauer et al., 2014). Hits were established by comparing each siRNA treated well with six control wells. We selected myosins for further analysis if average Golgi morphology of a siRNA treated well was significantly different to those of three independent control wells (50% of control) and were reproducible for one specific siRNA in three independent experiments (Figure 1B, red dots) or for two siRNAs in two independent experiments (Figure 1B, orange dots). This was the case for 17 myosins (about 50%), from seven different classes. We next tested the expression level of the identified myosins in hTertRPE-1 cells (Figure 1C). Skeletal and cardiac muscle myosins of the class II, expected not to be expressed in hTertRPE-1 cells, were expressed at the detection limit as judged by analysis of MYH6 and MYH15 (Figure 1C, note the logarithmic representation). Similarly, the expression levels of MYO3A, MYO10, MYO15A and MYH11 were at the detection limit, making a further analysis unfortunately very challenging. Yet, the mRNA of MYH9, MYO1C, MYO18A and MYO6 was well detectable (expression > 0.1% of GAPDH level). Probability density maps of the Golgi apparatus of pooled cells from significant wells revealed that gene silencing of MYO1C, MYO6 and MYO18 increased the area, in which Golgi structures were found as compared to pooled control cells (Figure 1D). Contrary, gene silencing of MYH9 decreased the area of the 50% Golgi apparatus probability contour (Figure 1D). MYH9, MYO18A and MYO6 have been all reported to regulate trafficking at the Golgi level (Billington et al., 2015; Müsch et al., 1997; Miserey-Lenkei et al., 2010; Warner et al., 2003) validating our density-based analysis. Additionally, our analysis identified MYO1C, as a potential novel regulator at the Golgi apparatus.

**Figure 1:**
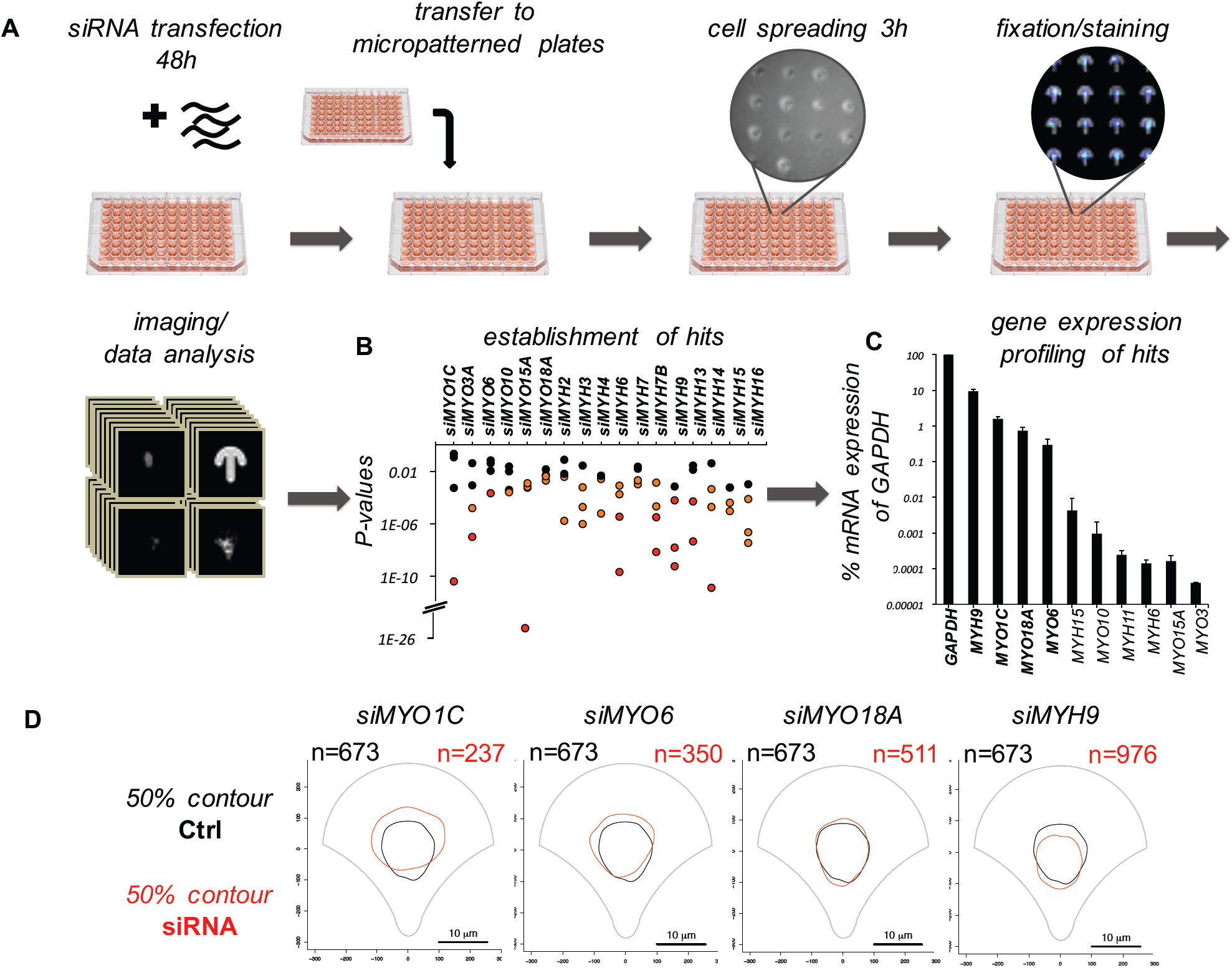
Identification of myosins that alter Golgi morphology. (**A**) Workflow for the systematic, comparative analysis combining cell culture on adhesive micropatterns and quantification of organelle positioning by probabilistic density mapping: human hTertRPE-1 cells were transfected with a siRNA library targeting each myosin with four independent siRNAs, incubated for 48h and transferred to crossbow-shaped micropatterned. After 3h spreading, cell were fixed, the Golgi apparatus was stained with GM130 antibody and imaged, following image processing and density-based statistical analysis. (**B**) A list of candidate myosins that were further investigated was established on the following criteria: average density map of the Golgi apparatus was significantly different to those of three independent control wells and were reproducible for one specific siRNA in tree independent experiments (red dots) or for two siRNAs in two independent experiments (orange dots). (**C**) The expression level of the identified myosins (excluding members of skeletal and cardiac muscle myosins) in hTertRPE-1 cells as a percentage of GAPDH expression. Error bars represent the standard deviation of three independent measurements. (**D**) Overlap of the 50% contour of the 2D density estimation map calculated in control cells (black) or cells depleted for the indicated myosin (red), n = number of analyzed cells.

### Comparable analysis of Golgi morphology upon depletion of different myosins

To further verify and characterize the identified myosins, we analyzed the 3D cellular phenotypes for individual siRNAs (Figure 2). The gene expression of all identified myosins significantly decreased to 10-40 % of control levels after siRNA induced gene silencing (Supplementary Figure 1A). Representative images of single cells (Figure 2A) and the volume of the 3D density maps of about 100 cells from three independent experiments confirmed that gene silencing of MYO1C, MYO6 and MYO18 increased the area, in which Golgi structures were found, as compared to pooled control cells, whereas gene silencing of MYH9 slightly decreased this area (Figure 2B, C). Moreover, automated detection of individual Golgi structures in single cells revealed that depletion of MYO1C and MYO18A significantly increased the average number of detectable Golgi structures per cell, indicating Golgi apparatus fragmentation (Figure 2D). Contrary, depletion of MYH9 significantly decreased the average number of Golgi structures per cell. Dividing the average numbers of structures per cell by the average volume of the 3D density map indicated that depletion of MYO1C and MYO6 de-compacted the Golgi apparatus by 40% and 25%, respectively. Depletion of MYO18A lead to a 50% compaction of this organelle whereas gene silencing of MYH9 did not change compaction of the Golgi apparatus to more than 10 % (Figure 2D). Thus, our results indicate that MYO1C and MYO6 depletion de-compacts the Golgi apparatus, whereas depletion of MYO18A and MYH9 leads to a more compact or smaller Golgi apparatus, increasing the number of Golgi structures in the case of MYO18A silencing and decreasing the Golgi area and number of Golgi structures downstream of MYH9 depletion.

**Figure 2:**
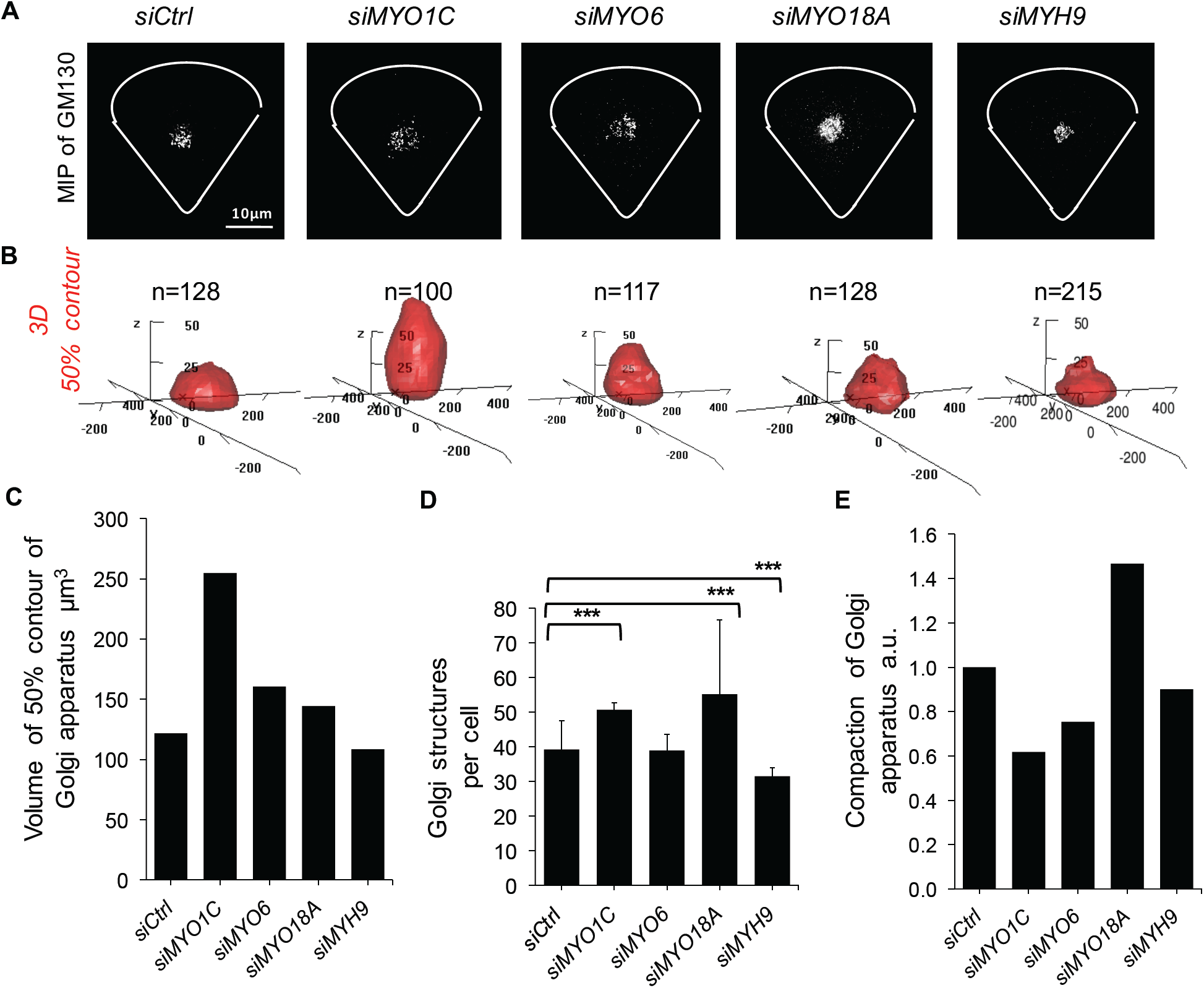
Comparable analysis of Golgi morphology upon depletion of different myosins. Fluorescent images representing the maximal intensity projections (MIP) of the Golgi apparatus stained by GM130 in single, representative hTertRPE-1 cells depleted for the indicated myosin and seeded on a crossbow-shaped micropattern. Scale bar: 10 µm. (**B**) 3D 50% density map of the Golgi apparatus as in **A**. n = number of analyzed cells. Axes are in pixels, 100 pixel = 6.45 µm. (**C**) Volume of the 50% contour of the Golgi apparatus as in **B**. (**D**) Quantification of the average numbers of Golgi structures per cell as in **B**. Error bars represent the standard deviation of three independent experiments. *** indicates P-value < 1×10^−4^ in a Student T-test. (**E**) Quantification of the compaction of the Golgi apparatus dividing average number of Golgi structures per cell through volume of 50% probability contour.

### Myosin 1C controls Golgi morphology

We next studied the function of MYO1C at the Golgi. As expected from mRNA levels, the MYO1C protein level was reduced to about 20% of control level after siRNA treatment (Supplementary Figure 1B, C). Analysis of the Golgi morphology in classical, unconstrained cell culture conditions confirmed that the Golgi apparatus was fragmented and the 2D projected Golgi area was significantly increased upon MYO1C depletion (Figure 3A, B). Notably, all tested Golgi marker, the matrix protein GM130, the *cis*-Golgi protein Giantin and the *trans*-Golgi network protein TGN46, exhibited altered morphology. Consistently, the overexpression of a GFP-MYO1C construct (of the rat homologue) in hTertRPE-1 cells let to a significant compaction of the Golgi apparatus that was not detected after the expression of GFP alone (Figure 3C, D). Overexpressed GFP-MYO1C showed similar distribution as the endogenous protein as quantified by average intensity projections of fluorescent images and density maps of endogenous and overexpressed MYO1C in micropatterned cells, and the average expression of EGFP-MYO1C protein level was comparable to endogenous levels (Supplementary Figure 1D-F). However, rescue experiments through the re-expression of this construct in MYO1C depleted cells were complicated by exhausted cell death after transfection. To study Golgi apparatus alterations at the ultrastructural level, we examined the MYO1C-depleted cells using electron microscopy. The EM showed that the characteristic structure of Golgi stacks, comprising four to six cisternae with a typical *cis-trans* Golgi and ER membrane, was detected after MYO1C depletion similar to the control cells, although the Golgi appeared less compact (Figure 3E).

**Figure 3:**
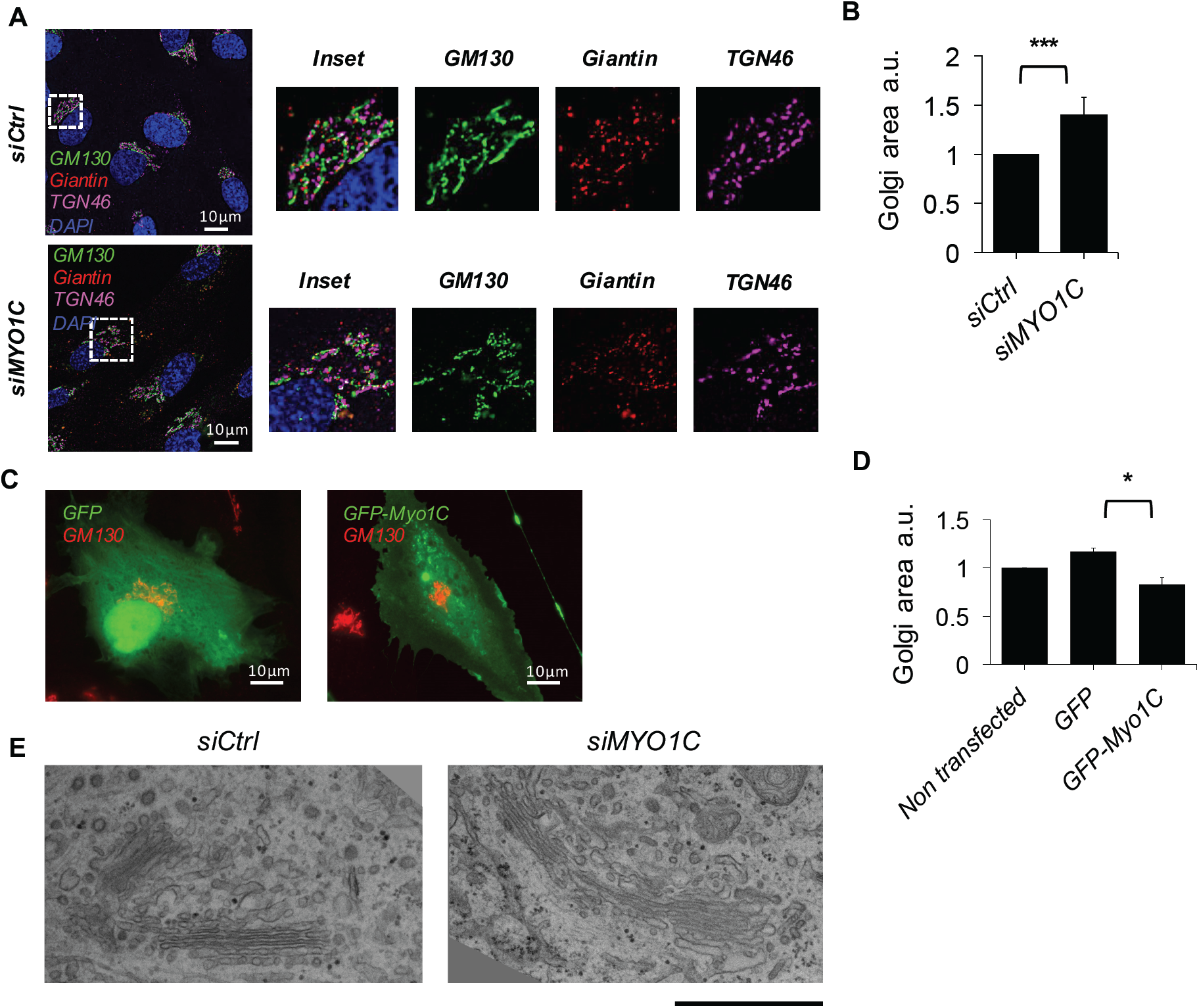
Myosin 1C controls Golgi morphology. (**A**) Immunofluorescent staining of different Golgi marker, the matrix proteins GM130 (green), the *cis*-Golgi protein Giantin (red) and the *trans*-Golgi network protein TGN46 (magenta) in representative control (top) and MYO1C depleted (bottom) hTertRPE-1 cells in classical, unconstrained culture conditions. The nucleus is stained with Dapi (blue). Quantification of the 2D projected Golgi area in control conditions (n=145) and upon MYO1C depletion (n=165). Error bars represent the standard deviation of three independent experiments. *** indicates P-value < 1×10^−4^ in a Student T-test. (**C**) Fluorescent images of representative hTertRPE-1 cells expressing GFP or GFP-MYO1C (green) and stained for the Golgi apparatus (red, GM130). (**D**) Quantification of the 2D projected Golgi area (stained by GM130) of non transfected cells and cells as in C. Error bars represent the standard deviation of three independent experiments with n > 40 cells of each condition. * indicates P-value < 1×10^−2^ in a Student T-test on averages of three independent experiments (**E**) Electron microscopy images of representative intracellular areas of unconstrained hTertRPE-1 cells containing the Golgi apparatus in control conditions and upon MYO1C depletion. Scale bar: 1 *µ*m.

### Myosin 1C accumulates at Golgi proximity by PH-tail domain and colocalizes with actin spots

The alteration in Golgi morphology after MYO1C depletion was somehow surprising, because MYO1C is known to be present at dynamic regions of the plasma membrane (Ruppert et al., 1995; Brandstaetter et al., 2012). Thus, we investigated whether there is an additional, intracellular pool of MYO1C. As expected, immunofluorescent staining of endogenous MYO1C or overexpression of EGFP-MYO1C indicated that the majority of MYO1C was at the plasma membrane. However, in addition to this major pool, we identified MYO1C-containing spots in close apposition to the Golgi apparatus (Figure 4A-C and Supplementary Figure 1G). Although MYO1C was in close contact with the *cis-, medial*- and *trans-* Golgi compartment, we could not detect significant colocalization with either Giantin, GM130 or TGN46. Live cell imaging of EGFP-MYO1C revealed that MYO1C-positive structures moved around the Golgi apparatus (Figure 4D Supplementary movie 1). Because MYO1C is an actin-based motor, we next analyzed whether MYO1C was found in actin-rich spots that are observed at the Golgi apparatus. Indeed, we found that both endogenous MYO1C and GFP-MYO1C colocalized with actin spots at the Golgi (Figure 5A, B). To verify that the recruitment of MYO1C at the proximity of the Golgi apparatus was independent of its binding to actin as previously reported (Tang and Ostap, 2001; Pyrpassopoulos et al., 2012), we compared accumulation of the overexpressed GFP-MYO1C with those of an actin-binding mutant GFP-MYO1CΔABL (Joensuu et al., 2014) in which the myopathy-loop (323–330; IIAKGEEL) was replaced by AGA tripeptide, disrupting the essential hydrophobic interactions between myosin head domain and actin filaments (Figure 5C). Additionally, we analyzed the accumulation of a truncated construct (GFP-Tail) containing only the MYO1C lipid-binding pleckstrin homology (PH) domain (Hokanson et al., 2006; Oh et al., 2013). We found that GFP-MYO1CΔABL formed the same numbers of spots at the Golgi as the non-mutated GFP-MYO1C indicating that accumulation of MYO1C at the proximity of the Golgi apparatus (marked by GM130) is independent of actin (Figure 5D, E). Overexpression of the GFP-Tail increased the presence of Golgi associated spots indicating that the PH-domain is responsible for accumulation around the Golgi. Interestingly, the expression of either GFP-MYO1CΔABL or the GFP-Tail did not change Golgi apparatus area (Supplementary Figure 2H). Together these results indicate that MYO1C accumulates at the proximity of the Golgi apparatus by its PH domain and that interaction of MYO1C with actin is important for Golgi morphology changes.

**Figure 4:**
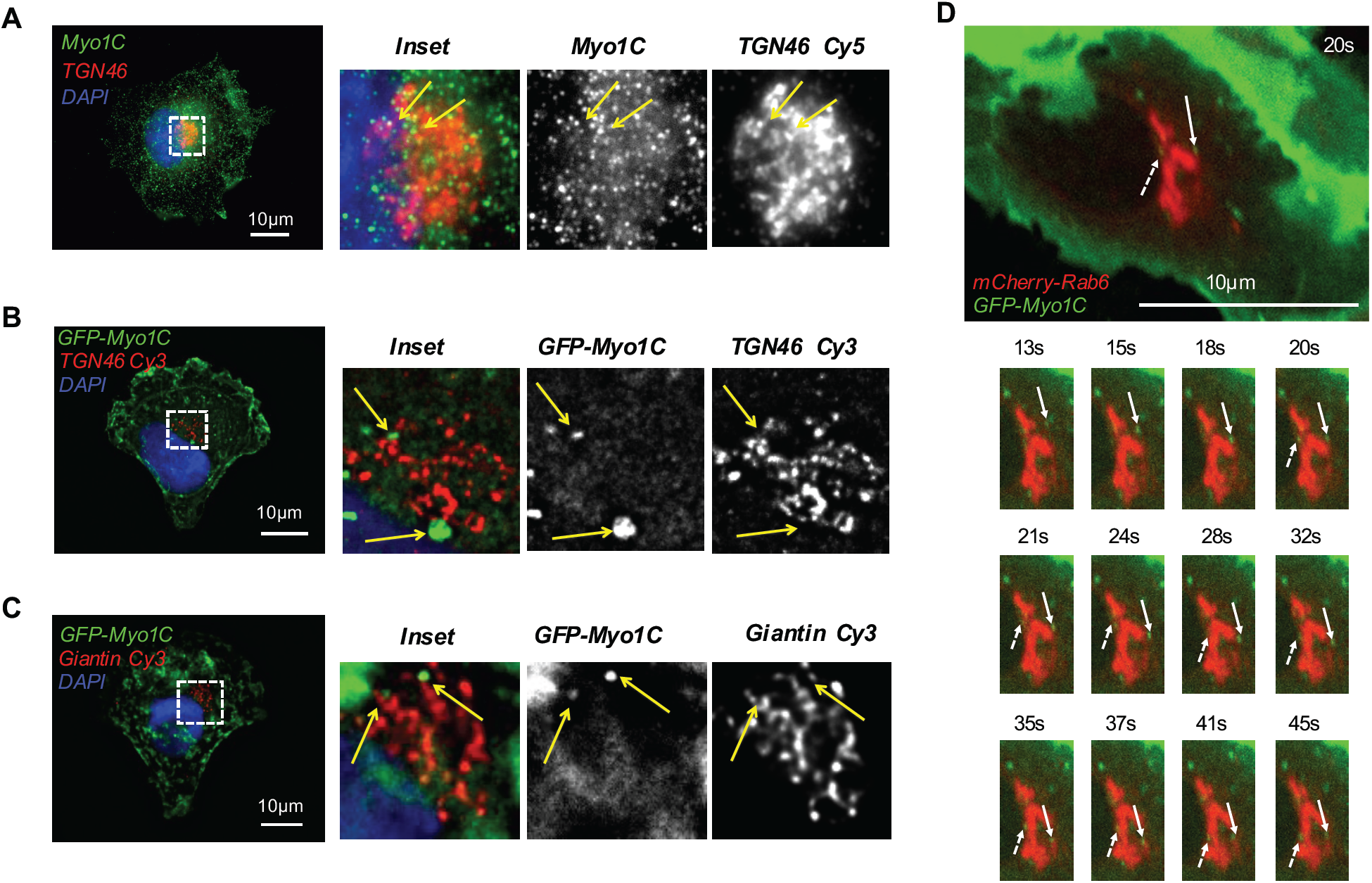
Myosin 1C accumulates at Golgi proximity. (**A**) Fluorescent images of a representative, unconstrained hTertRPE-1 cell stained with MYO1C antibody to visualize the endogenous protein (green), TGN46 antibody to visualize the Golgi apparatus (red) and Dapi to visualize the nucleus (blue). (**B, C**) Fluorescent images of representative, micropatterned hTertRPE-1 cells expressing GFP-MYO1C (green) and stained for the Golgi apparatus either with TGN46 or Giantin antibody (red) and the nucleus with Dapi (blue). (**D**) Fluorescent images of a time-lapse acquisition of unconstrained hTertRPE-1 cells expressing GFP-MYO1C (green) and mCherry-Rab6 (red). Scale bar: 10 µm.

**Figure 5:**
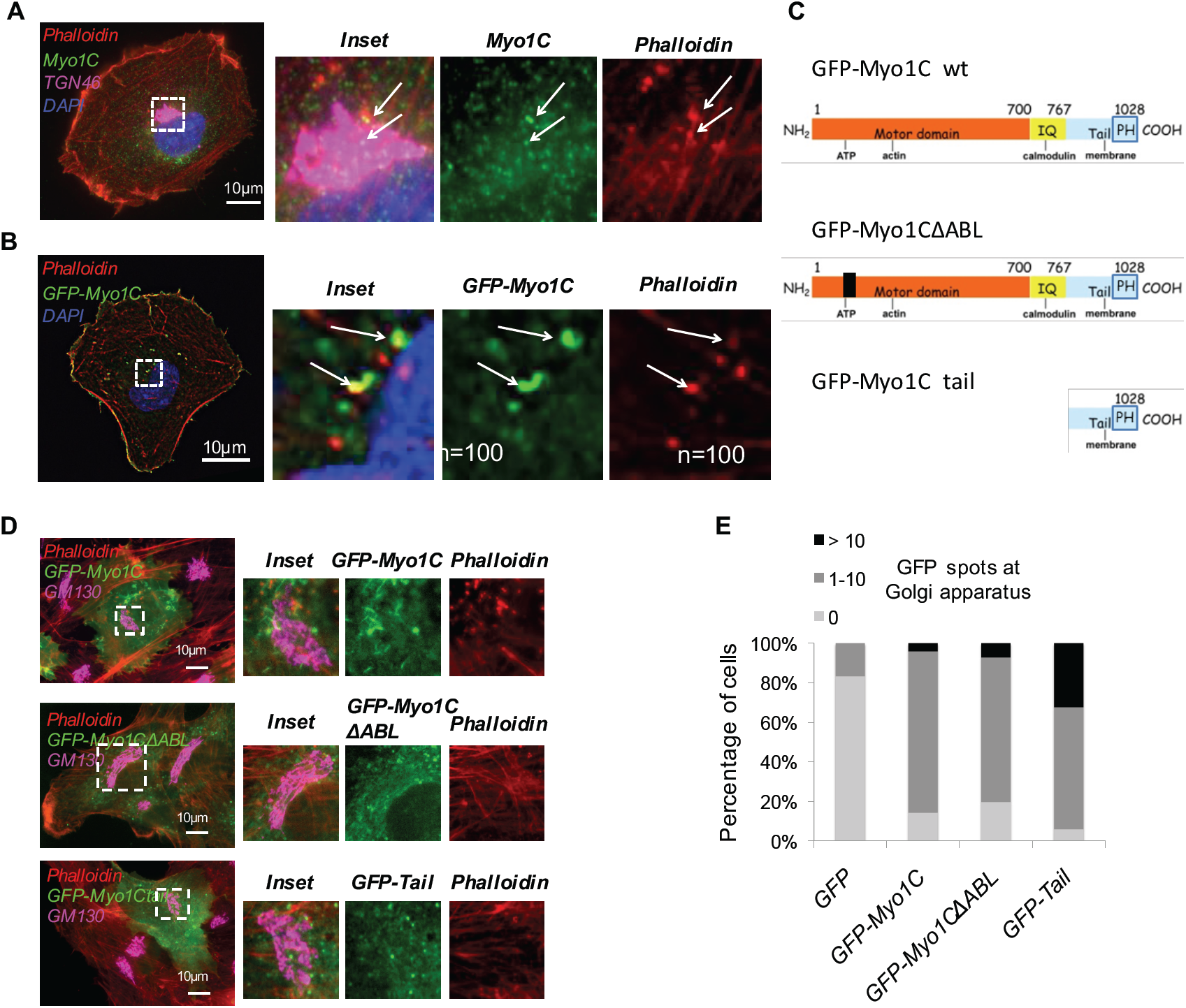
Myosin 1C colocalizes with actin at Golgi proximity but its recruitment is independent of its actin-binding domain. (**A**) Fluorescent images of a representative, unconstrained hTertRPE-1 cell stained with MYO1C antibody to visualize the endogenous protein (green), TGN46 antibody to visualize the Golgi apparatus (magenta) and phalloidin to visualize the actin cytoskeleton (red). The nucleus is stained with Dapi (blue). (**B**) Fluorescent images of representative, micropatterned hTertRPE-1 cells expressing GFP-MYO1C (green) and stained with phalloidin to visualize F-actin (red) and Dapi to visualize the nucleus (blue). (**C**) Domain structure of MYO1C, the mutant form MYO1CΔABL and the truncated Tail domain. (**D**) Fluorescent images of representative hTertRPE-1 cells expressing GFP-MYO1C, GFP-MYO1CΔABL or GFP-Tail (green), stained for the Golgi apparatus (magenta, GM130) and F-actin with phalloidin (red). (**E**) Quantification of GFP-positive spots at the Golgi area as in **D** for GFP (n=24), GFP-MYO1C (n=49), GFP-MYO1CΔABL (n=56) or GFP-Tail (n=34). % of cells is shown that contain 0 (light grey), 1-10 (dark grey) or more than 10 (black) Golgi-associated spots per cell. Note that control (GFP-expressing) cells never contained more than 1 Golgi-associated spot per cell.

### Myosin 1C controls cellular actin cytoskeleton by stabilizing branched actin

Next, we next investigated the organization of the actin cytoskeleton in the presence and absence of MYO1C employing micropattered, normalized cells for better comparison. Although actin protein levels were not changed upon MYO1C depletion (Figure 6A), staining of F-actin with phalloidin in normalized cells revealed a decrease of actin fibers throughout the cell upon MYO1C depletion (Figure 6B). This suggested that MYO1C stabilizes F-actin. Thus, we wondered whether actin depolymerization by different drugs could mimic MYO1C depletion. We treated cells with Latrunculin A to disrupt actin filaments (LatA, 200 nM for 20min) and CK666, an inhibitor of the Arp2/3 complex, to inhibit branched actin formation (CK666, 100 µM for 20 min) and observed the Golgi apparatus in hTertRPE-1 cells stably expressing the Golgi marker EGFP-Rab6A (Figure 6C). Whereas LatA treatment did not increase Golgi apparatus area, the loss of branched actin after inhibition of the Arp2/3 complex by CK666 significantly increased Golgi apparatus area (Figure 6D). Similarly, targeting ACTR3 of the Arp2/3 complex by siRNA resulted in a fragmented Golgi and an increase in Golgi area as compared to control in micropatterned cells (Figure 6E, F). These results indicated that MYO1C depletion mimics loss of branched actin. Based on our results, we propose that MYO1C stabilizes branched actin in the proximity of the Golgi apparatus.

**Figure 6:**
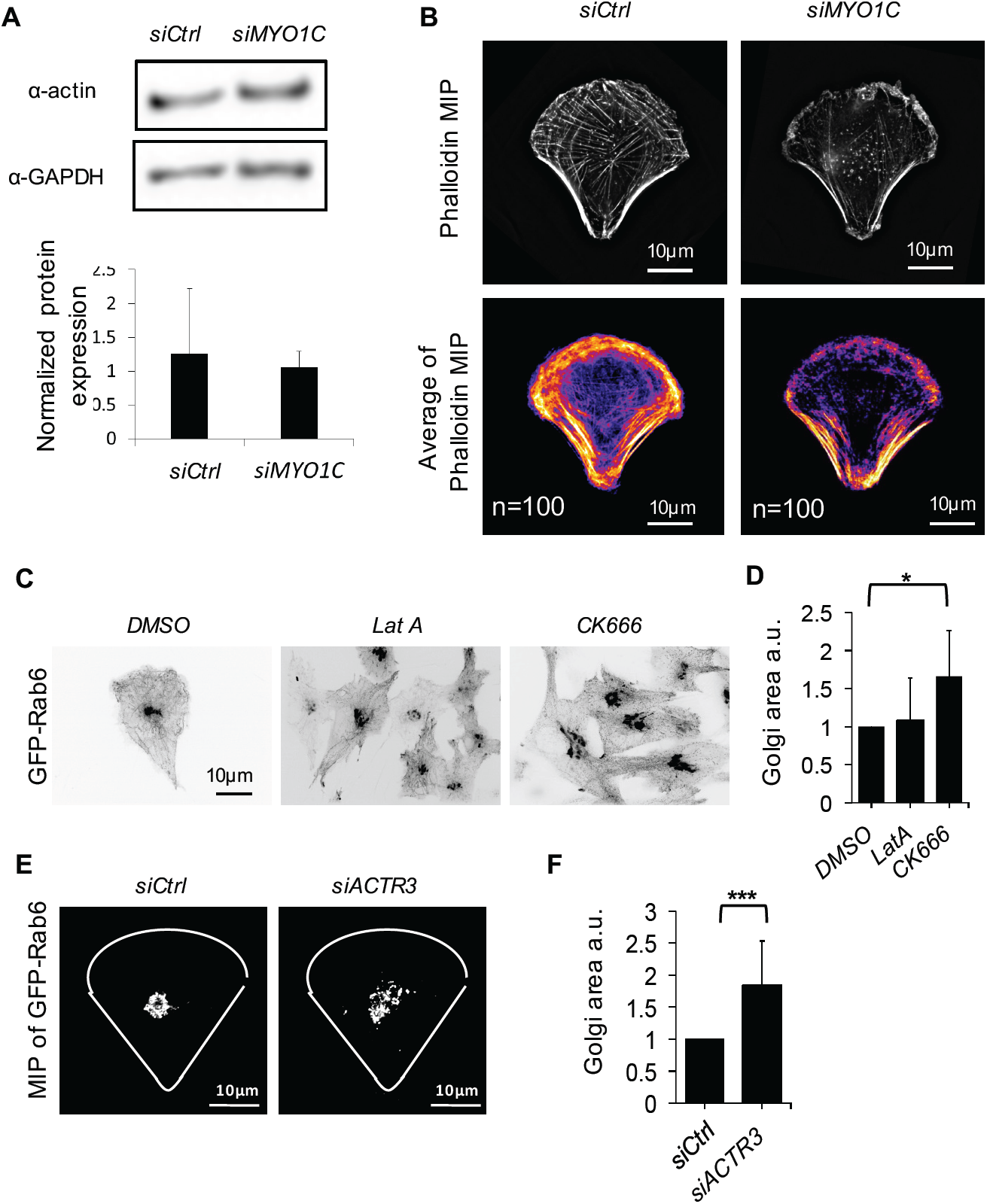
Myosin 1C controls cellular actin cytoskeleton by stabilizing branched actin. (**A**) Western blot analysis of actin and GAPDH expression levels in control and MYO1C depleted hTertRPE-1 cells (top) and densitometry quantification (bottom). Error bars represent the standard deviation of three independent experiments. (**B**) Fluorescent images representing the maximal intensity projections (MIP) of F-actin stained by phalloidin in representative control and MYO1C depleted hTertRPE-1 cells seeded on a crossbow-shaped micropattern (top). Average intensity projection of the MIP images from n cells from three independent experiments. Scale bar: 10 µm. (**C**) Fluorescent images of a representative, unconstrained hTertRPE-1 cells stably expressing GFP-Rab6 in control conditions and upon treatment with Latrunculin A and CK666. (**D**) Quantification of the 2D projected Golgi area (visualized by GFP-Rab6) in control conditions (n=14) and upon treatment with Latrunculin A (LatA, n=19) and CK666 (n=15). Error bars represent the standard deviation of individual cells. * indicates P-value < 1×10^−2^ in a Student T-test. (**E**) Fluorescent images representing the maximal intensity projections (MIP) of the Golgi apparatus (visualized by GFP-Rab6) in single, representative hTertRPE-1 cells in control conditions and ACTR3 depletion on a crossbow-shaped micropattern. Scale bar: 10 µm. (**F**) Quantification of the 2D projected Golgi area (visualized by GFP-Rab6) in control conditions (n=25) and ACTR3 depleted hTertRPE-1 cells (n=26). Error bars represent the standard deviation of individual cells. *** indicates P-value < 1×10^−4^ in a Student T-test.

### Myosin 1C delays arrival from anterograde and retrograde trafficking at the Golgi apparatus

To study the functional consequences of MYO1C depletion, we next investigated trafficking through the Golgi apparatus by live cell imaging. Employing the <u>r</u>etention of different reporter proteins in the ER <u>u</u>sing <u>s</u>elective <u>h</u>ooks of the RUSH system (Boncompain and Perez, 2012), we monitored synchronized anterograde transport of EGFP-TNFα from the ER to the plasma membrane after release of the reporter from the hook. As previously reported, EGFP-TNFα accumulated at the Golgi between 10 and 15 minutes (Boncompain and Perez, 2012) that we measured as increase in the ratio between mean intensity of EGFP-TNFα in GM130-positive Golgi area and a random cytosolic area of the same size (Figure 7A, B). Depletion of MYO1C let to a significant delay in the arrival at the Golgi apparatus at 10 min post pulse. In contrast, no difference between control and siMYO1C were detected after 35 minutes post pulse, suggesting that the exit of EGFP-TNFα from the Golgi is not delayed upon MYO1C depletion. We additionally analyzed a second cargo, EGFP-CD59, whose transport through the Golgi follows slower kinetics than those of TNFα: EGFP-CD59 passed the Golgi between 20-45 min post pulse. Consistently, we found that depletion of MYO1C significantly delayed arrival of EGFP-CD59 at the Golgi, accumulating there only 45 minutes post pulse (Supplementary Figure 2A, B). The Golgi apparatus can receive cargo from the retrograde pathway (Mallard et al., 1998). Thus, we next tested accumulation of Shiga toxin subunit B at the Golgi during its trafficking from preloaded endosomes to the ER according to standard protocols. STxB accumulated in the Golgi apparatus starting 15 min post release from endosomes, as previously reported. Again, we observed a significant delay in Golgi arrival upon MYO1C depletion (Figure 4C, D) at 15 and 90 min post release from endosomes. At 240 min post release, no differences between control and siMYO1C were detected indicating that the exit of STxB from the Golgi was not delayed upon MYO1C depletion. Thus, MYO1C facilitates the arrival at the Golgi from either anterograde or retrograde routes.

**Figure 7:**
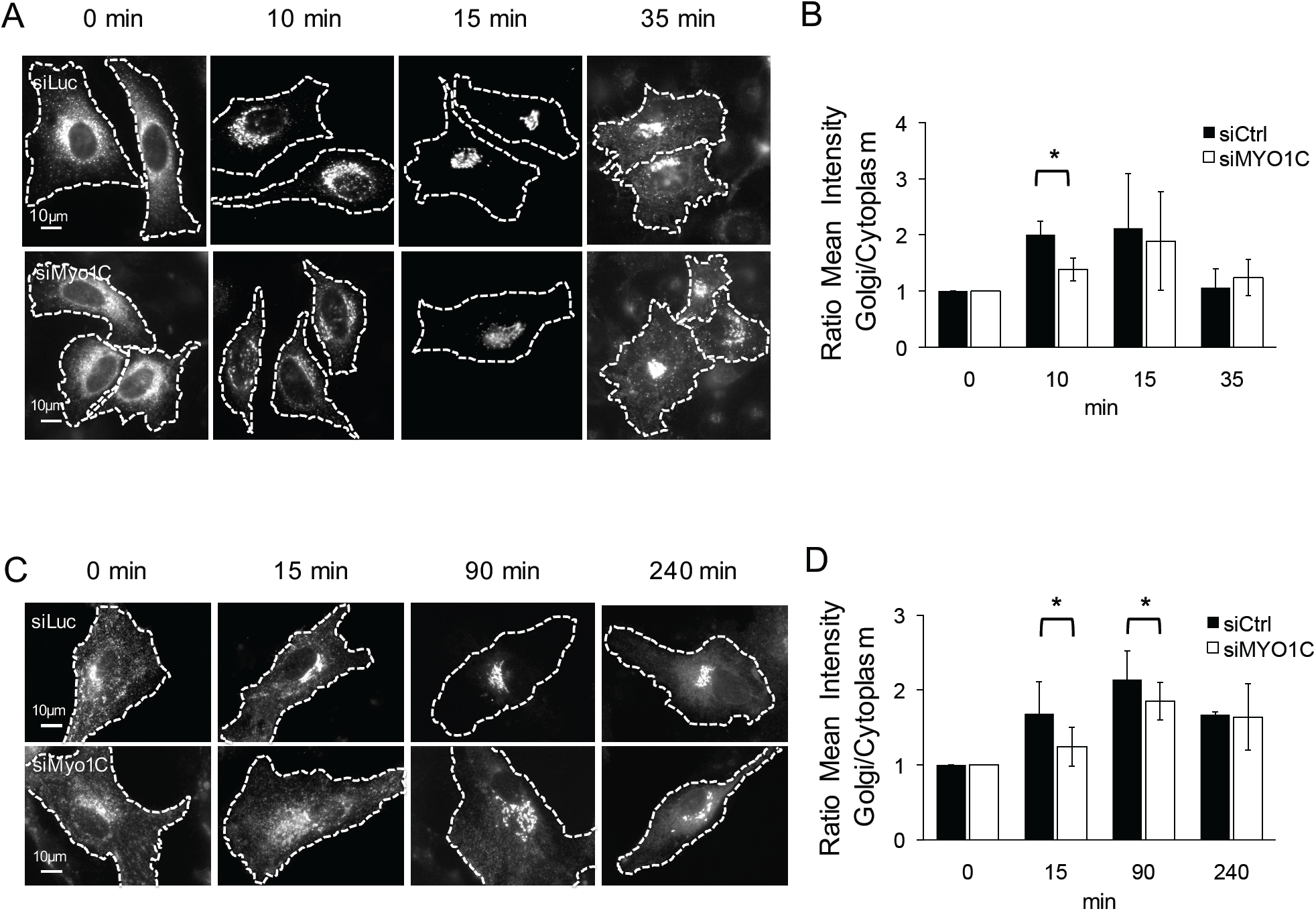
Myosin 1C delays arrival from anterograde and retrograde trafficking at the Golgi apparatus. (**A**) Fluorescent images of representative, unconstrained hTertRPE-1 cell stably expressing Str–KDEL_TNFα–SBP–EGFP at the indicated time points after the release of TNFα–SBP–EGFP from the ER in control conditions and in siMYO1C treated cells. Scale bar: 10 µm. (**B**) Quantification of the mean fluorescent intensity at the Golgi apparatus relative to a non-Golgi area of the same size as for A. Error bars represent the standard deviation of three independent experiment. * indicates P-value < 5×10^−2^ in a Student T-test on averages of three independent experiments. (**C**) Fluorescent images of representative, unconstrained hTertRPE-1 cell loaded with fluorescently labeled STxB at the indicated time points after the release from pre-loaded endosomes in control conditions and in siMYO1C treated cells. Scale bar: 10 µm. (**D**) Quantification of the mean fluorescent intensity at the Golgi apparatus relative to a non-Golgi area of the same size as for C. Error bars represent the standard deviation of three independent experiment. * indicates P-value < 5 × 10^−2^ in a Student T-test.

## Discussion

Based on our results we propose that MYO1C localizes at the proximity of the Golgi apparatus by its PH domain and stabilizes branched actin at the Golgi that facilitates arrival of incoming transport carriers at the Golgi apparatus (Figure 8). This proposed function of MYO1C differs substantially from those described for other Golgi-associated myosins, which have all been implicated in cargo exit from the Golgi apparatus. Indeed, myosins are required for the production of tensile forces for deformation of Golgi apparatus membranes for efficient tubule and vesicle formation and fission. Our work proposes that MYO1C is required for incoming Golgi cargos, organizing a scaffold of actin to facilitate arrival of cargos at membranes. The best-studied function of MYO1C is its role in the docking of GLUT4-containing vesicles to the plasma membrane in response to insulin stimulation (Boguslavsky et al., 2012; Bose et al., 2002, 2004; Chen et al., 2007; Huang et al., 2005; Yip et al., 2008). More generally, MYO1C has been shown to regulate exocytosis in secretory cells: MYO1C has been implicated in the VEGF-induced delivery of VEGFR2 to the cell plasma membrane (Tiwari et al., 2013) and in surfactant exocytosis (Kittelberger et al., 2016). Considering these studies and our novel results on the function of MYO1C at the Golgi, stabilizing branched actin for the arrival of transport carriers could be a general function of MYO1C in intracellular trafficking. We evidence that MYO1C depletion mimics the loss of the Arp2/3 complex at the Golgi apparatus. MYO1C has been previously shown to alter actin architecture in neuronal growth cones and at the immunological synapse in B cells (Joensuu et al., 2014; Maravillas-Montero et al., 2011). Interestingly, it has been shown that manipulation of the expression of another class I myosin, MYO1B, affect the distribution of Arp2/3 and associated branched F-actin (Almeida et al., 2011). Moreover, it has been shown in biochemical assays using geometrical patterning of actin that MYH9 and MYO6 can reorganize F-actin network architecture (Reymann et al., 2012). Thus, we propose that MYO1C reorganizes actin architecture. In the future, it will be important to address how MYO1C molecularly stabilizes branched actin. So far, it is unclear whether myosin-I motors influence actin dynamics by direct binding or by recruitment of factors that are involved in nucleation, polymerization and stabilization of actin filaments (McIntosh and Ostap, 2016; McIntosh et al., 2018; Evangelista et al., 2000).

**Figure 8:**
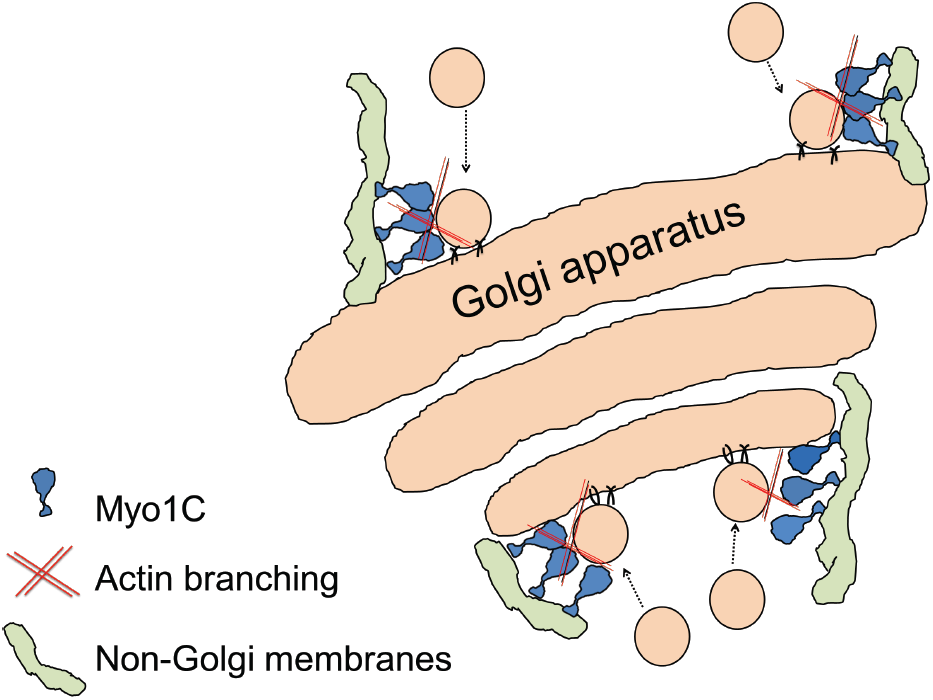
Model of MYO1C function at the Golgi apparatus. We propose that MYO1C accumulates at the proximity of the Golgi apparatus, on non-Golgi membranes, by its PH domain and stabilizes branched actin at the Golgi that facilitates arrival of incoming transport carriers at the Golgi apparatus.

In addition to MYO1C, we have identified MYO6, MYO18 and MYH9 (myosin IIA) as regulators of Golgi apparatus morphology in our density-based analysis. Although this confirms previous findings, the particular Golgi phenotype (for instance fragmentation, compaction) upon the depletion of a myosin seems to vary and to depend on several factors: i) cell spreading that can strongly impact the phenotypes of intracellular organelles, ii) cell type, reflecting variability in actin architectures, or iii) imaging techniques either visualizing a projected 2D area or the 3D volume of the Golgi apparatus. For instance, MYO18A depletion has been reported to prevent the lateral stretching of Golgi apparatus often observed in 2D culture (Dippold et al., 2010). Contrary, an expansion of the Golgi in the vertical direction as a result of actin filament disassembly downstream of MYO18A depletion was proposed by Bruun and colleagues (Bruun et al., 2017). Our approach on micropatterns has the advantage that it detects intracellular changes independently of changes in cell adhesion, and thus allows the comparably analysis after the depletion of different myosins in one cell type. Our results indicate that MYO1C and MYO6 depletion de-compacts the Golgi apparatus, whereas depletion of MYO18A and MYH9 leads to a more compact or smaller Golgi apparatus, increasing the number of Golgi structures in the case of MYO18A silencing and decreasing the Golgi area and number of Golgi structures downstream of MYH9 depletion. Quantitative comparative studies under normalized culture conditions are required in the future to comprehensively understand how the different myosins maintain the structural and functional integrity of the Golgi apparatus.

## Material and Methods

### Cells and reagents

Hunan telomerase-immortalized retinal pigmented epithelial (hTertRPE-1) cells (Invitrogen, Eugene, OR, USA) were grown in DMEM/F12 medium (Life Technologies, Carlsbad, CA, USA), supplemented with 10% Fetal Bovine Serum (FBS; Eurobio, Courtaboeuf, France), and 1% Penicillin-Streptomycin (Life Technologies), in a humidified atmosphere containing 5% CO_2_. The RUSH stable cell lines, either expressing Strep-KDEL_CD59–SBP–EGFP in hTertRPE-1 cells or Str–KDEL_TNFα–SBP–EGFP in HeLa cells, were kindly provided by Frank Perez (Institut Curie, France). For micropatterning experiments, fibronectin was from Sigma-Aldrich (St. Louis, MO, USA), fibrinogen–Cy5 from Invitrogen, and Poly-L-Lysine(20)-grafted[3.5]-Polyethyleneglycol(2) (PLL-g-PEG) from SuSoS (Dübendorf, Switzerland). The PLL-g-PEG was used at a final concentration of 0.1 mg.mL^−1^ in 10 mM HEPES (pH 7,3) solution. We used the following mouse monoclonal antibodies α-GM130 (BD Biosciences cat: 610823), α-MYO1C (Santa Cruz sc-136544) and α-actin (Sigma, A-2228), rabbit polyclonal antibodies α-Lamp1 (Sigma, L1418 or Abcam, Ab24170), human monoclonal antibody (F2C-hFc) α-tubulin and α-Giantin (Recombinant Protein and Antibody Platform of the Institut Curie) and sheep polyclonal antibody α-TGN46 (Bio-Rad, AHP500). FluoProbes 547H (557/572nm) coupled Phalloïdin was from Interchim. Cy3-coupled Shiga Toxin Subunit B (STxB) was generously provided by Ludger Johannes. Nuclei were marked using 0.2 μg.ml^−1^ 4’,6-diamidino-2-phenylindole (DAPI; Sigma-Aldrich).

### Cell transfection

Cells (200 000) were transfected in either 6 well plates or 12 well plates with 25 pmol.mL^−1^ siRNA (Sigma-Aldrich) using Lipofectamine RNAi MAX Transfection Reagent (5 μL.mL^−1^; Life Technologies). Cells were incubated 72 h prior further manipulations. MYO1C-siRNA 5’-GCUCAAAGAAUCCCAUUAU-3’; MYH9-siRNA 5’-GCACAGAGCUGGCCGACAA-3’; MYO18A-siRNA 5’-GGAGAUUAAUGGGCACAAU-3’; MYO6-siRNA 5’-CAAAGUCUGUUACUGAUUA-3’; ACTR3-siRNA CCGGCTGAAATTAAGTGAGGA were used.

Efficiency of siRNA gene silencing was verified by performing real time PCR on cell mRNA or Western Blot on cell lysates after three days of transfection. Controls were performed with siRNA targeting Luciferase. DNA transfection was performed using X-tremeGENE9 (Roche) following the manufacturer’s instructions. The plasmids used were GFP-MYO1C from Ana-Maria Lennon, GFP-MYO1C δABL from Eija Jokitalo, MYO1C tail from Woo Keun Song, mcherry-RAB6, mCherry-Rab6. The siRNA library targeting 36 myosins was kindly provided by Buzz Baum(Rohn et al., 2014).

### Screening procedure and analysis

A siRNA library targeting 36 myosin proteins (4 siRNAs/gene; 10 nM) was kindly provided by Baum Buzz (UCL). For siRNA transfection, hTertRPE-1 cells were seeded on black clear bottom 96-well plates (Perkin Elmer ViewPlate-96 Black, ref 6005182) at 70% confluency and transfected using 0.3 µl/well INTERFERin (Polyplus Transfection). Negative siRNA control (Luciferase, GL2) and a positive lethal siRNA (KIF11) were used as transfection controls. After 48 h, decreased cell viability in the lethal positive control wells served as an indicator of successful transfection (beta-factor was over 4 for each individual triplicate). Cells were then dissociated using Accumax™ solution (Sigma) and transferred to micropatterned-printed 96-well plates (CYTOO, Grenoble, France) containing crossbow micropatterns at a concentration of 4,000 cells/well. Plates were kept at 37°C for 3h and further cells were fixed with 4% (w/v) formaldehyde for 15 min and processed for immunofluorescence staining with α-GM130 primary antibodies. After washing three times with PBS, cells were stained with Cy3 donkey anti-mouse (Interchim, 1/200) in PBS/0.2% BSA/0.05% saponin for 1h at 37°C. Nuclei were stained with 0.2 μg/ml DAPI (Sigma). Three independent experiments were performed. Images were acquired using the INCell2000 automated imager (GE Healthcare). Acquisition was performed using a 20x dry objective and 42 fields per well were acquired giving rise to about 150 single cells per well. Images were sorted in ImageJ using the MSCS (Micropatterned Single Cell Sorting) plugin in ImageJ as previously described(Grossier et al., 2014a). Images of single micropatterned cells were segmented with MIA and the positional information of the Golgi apparatus was employed for density-based statistical analysis. Each of the wells containing cells transfected with siRNA against myosins was compared to six control wells on the same plate. The pairwise comparison of endosomal spatial distributions was carried out using the two-sample kernel density based test(Duong et al., 2012; Schauer et al., 2014). Conditions were considered as hits if average Golgi morphology of a siRNA treated well was significantly different (*P*-value < 0.0005) to those of three independent control wells and were reproducible for one specific siRNA in tree independent experiments or for two siRNAs in two independent experiments.

### Micropatterned coverslips preparation and cell seeding

Micropattern production was as previously described(Azioune et al., 2009) using photo-lithography methods. Briefly, coverslips were coated with PLL-g-PEG and spatially controlled areas were exposed to deep UV during 5 min using a photomask. Crossbows (37 μm diameter, 7 μm thick) were therefore photo-printed. Prior to cell seeding, the patterned substrates were incubated for 1h with fibronectin at a concentration of 50 mg.mL^−1^. The fibronectin mixture was supplemented with 10 mg.mL^−1^ fibrinogen– Cy5 to stain the micropatterns. Cells were seeded on micropatterns in DMEM/F12 medium supplemented with 10 mM HEPES for 3 h prior the experiment.

### Drug treatments

To induce cytoskeleton changes, the following drugs were used at the indicated concentrations for 20 min: Latrunculin A 200 nM (Sigma-Aldrich, L5163), CK666 100 *µ*M (Sigma-Aldrich, SML0006). DMSO was used for control conditions.

### Immunofluorescence staining

For immunofluorescence staining, formaldehyde-fixed cells were washed three times with PBS and permeabilized in PBS/0.2% BSA/0.05% saponin. Cells were then incubated with a primary antibody for 1 h, washed in PBS and incubated with Alexa Fluor 488- or Cy3-coupled secondary antibodies (Jackson ImmunoResearch). Slices were mounted in Mowiol (Sigma-Aldrich).

### Synchronization of protein secretion by RUSH system

Stable cell lines expressing Strep-KDEL_CD59–SBP–EGFP (hTertRPE-1 cells) and Str–KDEL_TNFα– SBP–EGFP (HeLa cells) were transfected with MYO1C-siRNA or Luciferase-siRNA. After 72 h, cells were treated with 40 μM biotin to allow the reporter release into the secretory pathway. Cells were fixed at the indicated time points after biotin addition, stained with against GM130 and imaged.

### STxB assay

hTertRPE-1 cells were transfected with MYO1C-siRNA or Luciferase-siRNA. After 72 hours, cells were incubated with Cy3-STxB (1μg/ml) for 1 hour at 19°C to load STxB in early and recycling endosomes followed by three washed with PBS. Fresh medium was added and cells were transferred at 37°C for 15, 90 and 240 min, fixed, stained with against GM130 and imaged.

### Electron microscopy (EM)

hTertRPE-1 cells were transfected with siRNA-MYO1C or siRNA-LUC for 72 hours and chemically fixed with 2% glutaraldehyde and 1% paraformaldehyde in NaHCa buffer (100 mM NaCl, 30 mM HEPES, and 2 mM CaCl_2_ at pH 7.4). The specimens were postfixed with 1% OsO4/1.5% K4Fe(CN)6, dehydrated with a graded ethanol series, and embedded in Epon 812 (TAAB Laboratories Equipment Ltd.). Ultrathin sections were mounted on EM grids, and stained with uranylacetate and lead citrate. Sections were observed with a transmission electron microscope (Tecnai Spirit, Thermo Fisher Scientific, Eindhoven Netherlands) and digital acquisitions were made with a numeric camera (Quemesa; EMSIS, Münster, Germany).

### Fluorescent image acquisition

For micropatterned cells, Z-stack images from fixed and immunolabelled cells were acquired with an inverted widefield Deltavision Core Microscope (Applied Precision) equipped with highly sensitive cooled interlined chargecoupled device (CCD) camera (CoolSnap Hq2, Photometrics). Z-dimension series were acquired every 0.2 *µ*m and out of focus signals were reduced using deconvolution. Non-constrained cells were examined using a three-dimensional deconvolution microscope (Leica DM-RXA2), equipped with z-drive (Physik Instrument) and 100 × 1.4NA-PL-APO objective lens. Three-dimensional or one-dimensional multicolor image stacks were acquired using the Metamorph software (MDS) through a cooled CCD camera (Photometrics Coolsnap HQ). Z-dimension series were acquired every 0.2 µm. For video microscopy experiments, transfected cells were grown on glass bottom Fluorodishes. Video microscopy was performed at 37 °C using a spinning-disk microscope mounted on an inverted motorized microscope (Nikon TE2000-U) through a 100 × 1.4NA PL-APO objective lens. The microscope was equipped with a Yokogawa CSU-22 spinning-disk head, a Roper Scientific laser launch, a Photometrics Coolsnap HQ2 CCD camera for image acquisition and Metamorph software (MDS) to control the setup. Z-dimension series were acquired every 0.2 µm.

### Image analysis

Golgi area was selected based on GM130 staining and the area and mean intensity were measured in ImageJ(Rasband). Values were normalized to control conditions (non transfected, DMSO or siLUC). In anterograde and retrograde transport assays a small area in the Golgi (GM130-positive) and a random cytoplasmic area of the same size was selected and mean fluorescent intensities were measured for GFP-TNFα, GFP-CD59, Cy3-ShTx using ImageJ. Values were expressed as mean intensity ratio Golgi/Cytoplasm.

### Statistical analysis

For each experiment, a large number of cells were monitored from 3 independent experiments. Data from independent experiments were combined for representation and statistical analysis was performed employing a bilateral Student t-test.

### Kernel density estimation

To extract the 3D spatial coordinates of intracellular structures, images of several tens of cells were segmented with the multidimensional image analysis (MIA) interface on MetaMorph(Molecular Devices) based on wavelet decomposition. The coordinates from all cells in one condition were aligned according to the micropattern as previously described(Grossier et al., 2014b; Schauer et al., 2010). The aligned coordinates of the segmented structures were processed for density estimation programmed in the ks library in the R programming language(R Development Core Team, 2013): the probability density function *f* for each data sample of *n* coordinates *X*_1_, *X*_2_, *…, X*_n_ was estimated. We used a non-parametric, unbinned kernel density estimator. At each of the data points, a kernel function *K* was centered. The kernel functions were then summed to form the kernel density estimator 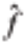

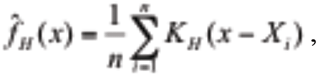

in which k_H_ is the Gaussian kernel with mean zero and variance matrix *H*. To estimate *H* (also known as the bandwidth), we used the plug-in selector in the ks library that has been shown to be reliable for 2D and 3D spatial distributions. For visualizing kernel density estimates, we used probability contours and the extension libraries mvtnorm, rgl, and miscd.

## Acknowledgements

We acknowledge the following people for providing materials: Franck Perez, Ana-Maria Lennon-Duménil, Eija Jokitalo and Woo Keun Song for the plasmids. Buzz Baum for the myosin library. We thank Tarn Duong for advices on statistical analysis and kernel density estimation and Sarah Tessier for the siRNA screening technical assistance. The authors greatly acknowledge the Cell and Tissue Imaging Facility (PICT-IBiSA @Burg, PICT-EM @Burg and PICT-IBiSA @Pasteur) and Nikon Imaging Center, Institut Curie (Paris), member of the French National Research Infrastructure France-BioImaging (ANR10-INBS-04). This work was supported by the Labex CelTisPhyBio (ANR-10-LBX-0038) and Idex Paris Sciences et Lettres (ANR-10-IDEX-0001-02 PSL). This project was supported by grants from Agence Nationale de la Recherche (#2010 BLAN 122902), European Research Council advanced grants (project 339847 ‘MYODYN’), the Centre National de la Recherche Scientifique and Institut Curie.

